# Controlling wireframe DNA origami nuclease degradation with minor groove binders

**DOI:** 10.1101/2020.05.24.110783

**Authors:** Eike-Christian Wamhoff, Hellen Huang, Benjamin J. Read, Eric Ginsburg, William R. Schief, Nicholas Farrell, Darrell J. Irvine, Mark Bathe

## Abstract

Virus-like DNA nanoparticles have emerged as promising vaccine and gene delivery platforms due to their programmable nature that offers independent control over size, shape, and functionalization. However, as biodegradable materials, their utility for specific therapeutic indications depends on their structural integrity during biodistribution to efficiently target cells, tissues, or organs. Here, we explore reversible minor groove binders to control the degradation half-lives of wireframe DNA origami. Bare, two-helix DNA nanoparticles were found to be stable under typical cell culture conditions in presence of bovine serum, yet they remain susceptible to endonucleases, specifically DNAse I. Moreover, they degrade rapidly in mouse serum, suggesting species-specific degradation. Blocking minor groove accessibility with diamidines resulted in substantial protection against endonucleases, specifically DNAse-I. This strategy was found to be compatible with both varying wireframe DNA origami architectures and functionalization with protein antigens. Our stabilization strategy offers distinct physicochemical properties compared with established cationic polymer-based methods, with synergistic therapeutic potential for minor groove binder delivery for infectious diseases and cancer.

## Introduction

DNA nanotechnology offers tremendous potential for the development of next-generation vaccine and gene therapeutics^1-2^. Biodegradable DNA nanoparticles provide full control over their size and shape, as well as site-specific functionalization for active targeting or incorporation of gene therapeutic cargos and immunomodulatory adjuvants. In particular, the scaffolded DNA origami method produces monodisperse nanoparticles at the 10-200 nm scale and near quantitative yields^3^. The two major classes of DNA origami developed over the past decade are densely-packed, brick-like assemblies^4^ and wireframe assemblies^5^. The development of top-down sequence design algorithms has enabled facile prototyping of the latter class of objects^6-9^, and scalable enzymatic and bacterial production of single-stranded DNA scaffolds with custom sequence and length have paved the way for pre-clinical and clinical studies^10-13^. DNA nanostructures have further been deployed in a variety of preliminary studies as therapeutic delivery platforms. These applications include the controlled organization of antigens to activate B cells^14^, the delivery of siRNA^15-16^, CpG oligonucleotides^17^ and small molecules drugs^18-20^. Logic-gated nanorobots have been leveraged to to achieve controlled cargo release in vitro^21^ and in vivo^1^.

Unlike polymeric, liposomal, and protein-based nanoparticles, DNA-based materials are, however rapidly biodegradable due to nuclease activity^22-24^. In addition, multi-layer brick-like assemblies are subject to disassembly at physiological magnesium concentrations^25^, in contrast to DX-based, two-helix wireframe architectures (hereafter termed two-helix to contrast one-helix and six-helix assemblies)^7^. While their biodegradability is attractive from a toxicity and safety standpoint, nuclease degradation of native, unmodified DNA nanoparticles may limit their serum half-life to several minutes^23-24^, which is insufficient for vaccine or gene therapeutic applications that require lymph trafficking or penetration of other target tissues, which may require hours in larger mammals, particularly humans.

While the nuclease degradation of brick-like DNA origami has been studied extensively, finding they are largely exonuclease resistant and endonuclease susceptible^26^, virus-like wireframe assemblies composed of have only been examined to a limited extent. For the former class of objects, numerous stabilization strategies have been explored, including lipid encapsulation^23^, coating with cationic oligo- or polymers^24, 27-28^, PEGylation^29^ and combinations thereof. Recently, glutaraldehyde was employed as a crosslinker to irreversibly attach coating agents, providing exceptional endonuclease protection and degradation half-lives of several days^30^. Additionally, photoinduced cross-linking of thymidines^25^ or chemical ligation at nick positions^31^ to create topologically interlocked brick-like assemblies eliminated the requirement for high magnesium concentration to stabilize brick-like DNA origami from electrostatically-induced disassembly.

Yet, the preceding strategies display several potential disadvantages. For instance, lipid encapsulation is associated with challenges around solubility and dispersity. Cationic oligo- and polymers change the physicochemical properties of the DNA nanoparticle and thereby its biodistribution while PEGylation might limit the accessibility of targeting ligands or antigens. Further, chemical crosslinking might not be compatible with DNA origami functionalization. As each therapeutic application poses unique technological challenges, the development of alternative, complementary stabilization strategies remains of importance for the DNA nanotechnology field.

Because DNAse I represents the major extracellular endonuclease in mammals, found in blood and interstitial fluids, we sought to specifically inhibit its activity and ideally control the rate of degradation, accounting for species- and tissue-specific expression levels^32-34^. DNase I preferentially cleaves dsDNA over ssDNA, with slight preferences for AT tracts. It recognizes dsDNA via the minor groove with cleavage involving bending of the double-helix towards the major groove^35-36^. Accordingly, minor groove binders (MGBs) have previously been used to modulate DNAse I-dependent degradation of dsDNA, and are commonly used in footprinting assays^37-38^. Three major classes of canonical MGBs have been described: diamidines, benzimidazoles, and pyrrole-imidazole polyamides^39^. Extensive medicinal chemistry efforts have resulted in to the development of FDA-approved experimental drugs or clinical candidates for the treatment of both infectious diseases and cancer, with inhibition of replication being the primary mode of action (NCT 03824795 is an ongoing phase II clinical trials)^40-41^. MGBs typically display preferential binding to AT tracts, yet their specificity and affinity are readily tunable^42-43^. A related class of compounds, platin-based phosphate clamps (PCs, hereafter also termed MGBs), interact with the phosphate backbone but have also been shown to restrict access to the minor groove of dsDNA and to thereby modulate DNAse I activity in footprinting assays^44-46^. While binding of non-canonical MGBs to DNA origami has been characterized, protection against endonucleases was not evaluated^47^.

Here, we investigate the stabilization of wireframe DNA origami using MGBs **(Figure 1).** Characterization in cell culture conditions revealed long-term stability of bare two-helix wireframe architectures of more than 24 h. We further found that these DNA nanoparticles are exonuclease resistant but remain susceptible to endonucleases at physiological concentrations and notably when incubated in mouse serum compared with fetal bovine serum. To stabilize two-helix wireframe architectures, we explored compounds known to restrict access to the minor groove and to thereby competitively inhibit DNAse I activity. Toward this end, we compared representatives of two classes of canonical minor groove binders, diamidines and benzimidazoles, with platin-based PCs **(Figure 1)**. All compound classes increased degradation half-lives in mouse serum, with the diamidine **5** (2-(4-Amidinophenyl)-1H-indole-6-carboxamidine, DAPI) exhibiting the strongest stabilization effect. Finally, we found that diamidine-based protection from endonucleases was compatible with varying DNA nanoparticle shapes and virus-like DNA origami nanoparticles functionalized with the clinical HIV-1 vaccine candidate eODGT8. Hence, the use of MGBs represents a facile and generalizable stabilization strategy that paves the way for potential therapeutic applications. It may also offer the opportunity for the challenging delivery of these clinically relevant compounds to combat infectious diseases and cancer, depending on dosing requirements.

**Figure 1.**
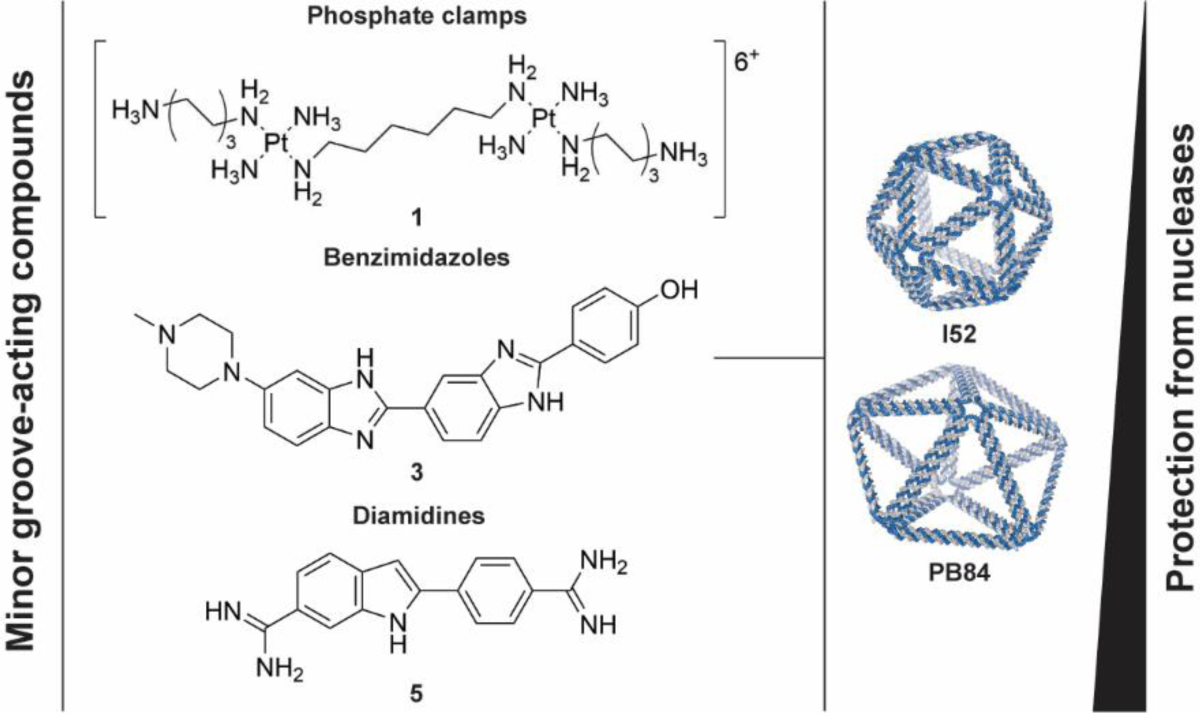
Stabilization of wireframe DNA origami with minor groove binders. Overview of screened classes of MGBs sorted by level stabilization against nuclease degradation conveyed to wireframe DNA origami. Diamidine **5** (DAPI) represents the most potent stabilizer identified and was found to be compatible with different geometries and edge architectures as well as DNA origami functionalization.

## Results and Discussion

We characterized the stability of a two-helix pentagonal bipyramid (**PB84**) designed with DAEDALUS^7^ under cell culture conditions **(Figure S1, Tables S1 to S3)**. Prompted by our previous findings that this class of wireframe assemblies is stable in PBS^7^, we envisioned that nanostructuring provides sufficient protection against nucleases for in vitro experiments. The DNA nanoparticle was first incubated in DMEM containing 10% fetal bovine serum (FBS), with observed half-lives beyond 24 h, whereas the unfolded, circular scaffold was substantially degraded within 3 h **(Figure 2a)**. Additional characterization via gel electrophoresis under denaturing conditions revealed that oligonucleotide staples remained intact up to 48 h, while the onset of DNA nanoparticle degradation appeared to coincide with nicking of the circular scaffold as early as 12 h **(Figure 2b**). In accordance with these observations, **PB84** was resistant to bacterial exonuclease Exo I degradation, but was rapidly degraded at physiological DNAse I concentrations **(Figure 2c)**. Notably, the protection of 3’ and 5’ termini of the oligonucleotide staples using hexaethylene glycol did not further stabilize **PB84** against 10% FBS **(Figure S2)**^29^.

**Figure 2.**
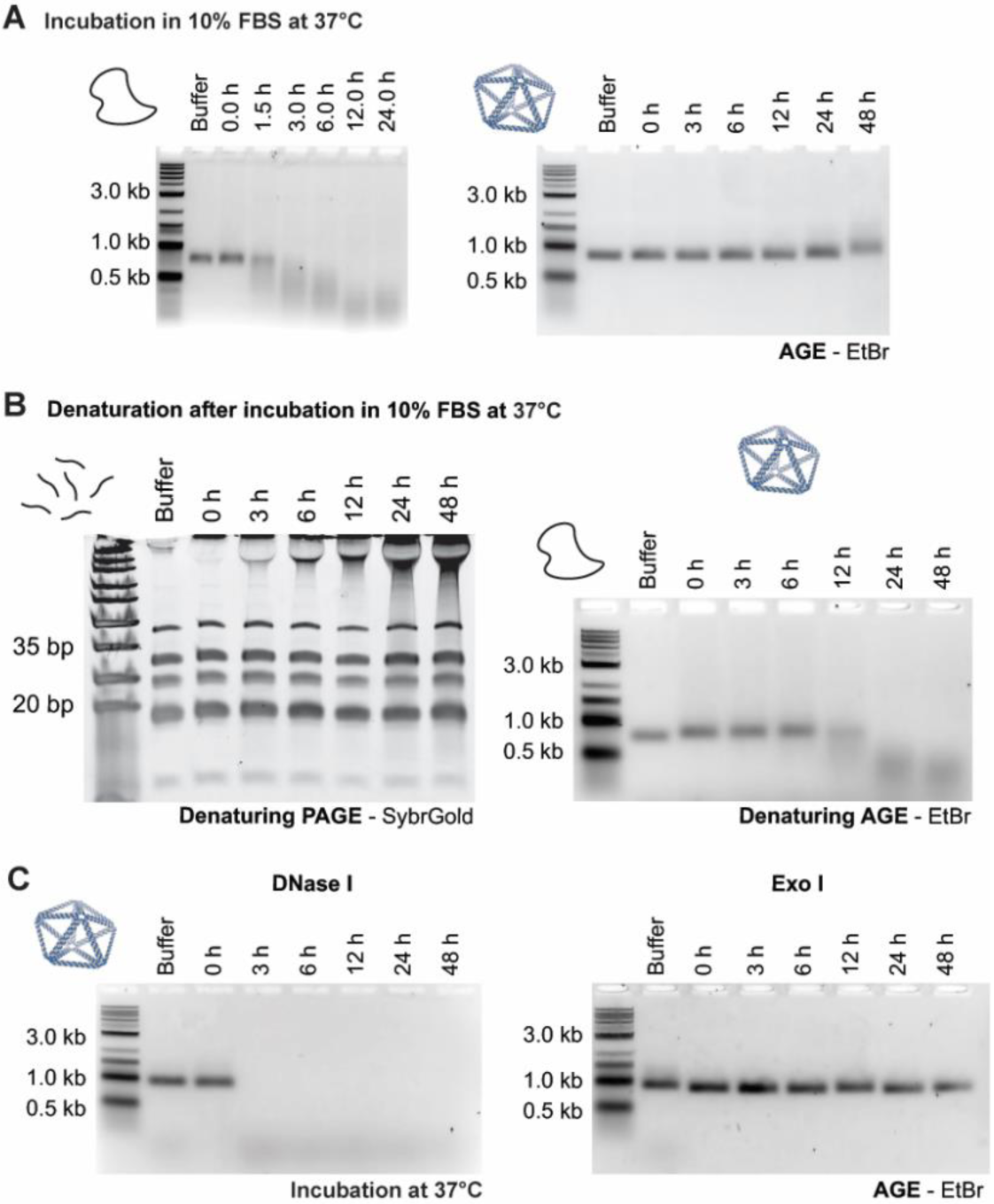
Stability of two-helix wireframe DNA origami under cell culture conditions. **(A)** Incubation under typical cell culture conditions, in DMEM with 10% FBS revealed stability of **PB84** for more than 24 h. The circular ssDNA scaffold fully degraded within 3 h under these conditions. **(B)** Gel electrophoresis under denaturing conditions indicates that the onset of **PB84** degradation coincides with nicking of the scaffold while staple oligonucleotides only show negligible degradation. **(C)** This observation is in accordance with the resistance of **PB84** against exonuclease Exo I and its susceptibility against DNase I at physiological concentrations (0.500 U/ml). All experiments were performed in triplicate. Representative gel images are shown.

We conclude that two-helix wireframe assemblies, exemplified by **PB84**, are stable under typical cell culture conditions, and resistant to exonuclease degradation. However, they remain susceptible to endonuclease activity, with DNAse I being the most abundant nuclease in FBS. We hypothesize the observed endonuclease preference for the scaffold over oligonucleotide staples to be due to a combination of three factors. First, band shifts are very sensitive for the introduction of nicks in circular ssDNA. Secondly, the likelihood of introducing a nick on the scaffold is higher overall than for any individual staple given its increased length. Finally, DNAse I activity depends on local double-helix flexibility and is thus expected to be increased at nick positions where it can only cleave the scaffold.

Next, we tested the stability of two-helix wireframe assemblies under conditions more closely resembling the in vivo extracellular environment of pre-clinical mouse models. Toward this end, we incubated **PB84** in 10% mouse serum (MS) **(Figure 3a, Figures S3)**. Strikingly, we observed rapid degradation of the DNA nanoparticle within 3 h, highlighting the importance of accounting for species-specific DNase-I activity for pre-clinical studies using DNA origami^32-34^. We further suspect that different DNAse I orthologs might display unique substrate specificity with regard to nanostructuring. Similar characteristics have previously been described for both murine and human DNAse I homologs in lupus erythematosus, where the degradation of chromatin is associated with pathogenesis^48-49^.

**Figure 3.**
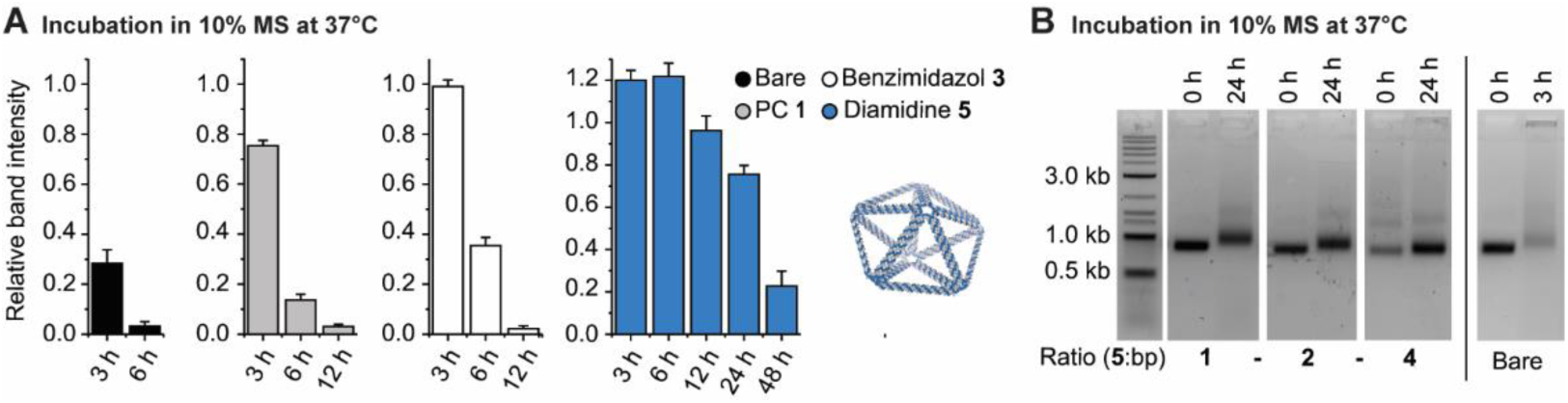
Stabilization of two-helix wireframe DNA origami with diamidines. **(A)** The screening of MGBs **1** to **5** revealed substantial, yet differential stabilization of **PB84** against 10% MS. Relative band intensities compared to the 0 h data point for bare and coated DNA nanoparticles are shown. Diamidine **5** was identified as the most potent stabilizer conveying half-lives beyond 24 h to **PB84. (B)** Increased concentrations of **5** further stabilize wireframe DNA origami, delaying band shifts diagnostic of the degradation onset. All experiments were performed in triplicate. Representative gel images are shown.

Compounds restricting access to the minor groove of dsDNA have been reported to competitively inhibit DNAse I activity, including two of the canonical MGB classes^37-38^, benzimidazoles and diamidines, as well as platin-based PCs^45^. We therefore screened representative compounds of all three MGB classes for their capacity to protect DNA nanoparticles against endonuclease degradation **(Figure 3a, Figure S3)**. Following incubation at MGB:base pair ratios of 1, all classes extended the half-life of **PB84** in 10% MS. Yet, we observed differential levels of endonuclease inhibition, with PC **1** providing protection for only up to 3 h. Incubation with PC **2**, in contrast, resulted in aggregation of **PB84 (Figure S3)**. Benzimidazole **3**, a commonly used Hoechst dye also characterized in clinical studies for the treatment of pancreatic carcinoma under the synonym pibenzimol, further extended the half-life of **PB84**^40^.

By comparison, the tested diamidines **4** and **5** conveyed superior stabilization against 10% MS. Incubation with **4**, also known as furamidine, an FDA-approved experimental drug for the treatment of pneumocystis pneumonia, extended the half-life of the DNA nanoparticle beyond 6 h **(Figure S4)**. Diamidine **5**, 2-(4-amidinophenyl)-1H-indole-6-carboxamidine (DAPI), represents the most potent stabilizer tested, providing protection for up to 24 h **(Figure 3a)**. Notably, the onset of degradation of **PB84** was characterized by subtle band shifts, followed by the rapid loss of total band intensity. Incubation at increased MGB:base pair ratios mitigated this band shift for up to 24 h and further improved protection by **5 (Figure 3b)**. Incubation at a ratio of 4 resulted in reduced band intensities for the 0 h time point (also observed for ratios of 1 and 2, albeit to a lesser extend) and partial aggregation of DNA nanoparticles. We suspect that this is caused by the previously reported secondary, intercalating binding mode of **5**^37^.

In summary, these findings demonstrate that MGBs protect two-helix wireframe DNA origami degradation by DNAse I, with the degree of protection depending on the compound class. Diamidine **5** was identified as the most potent stabilizer that increased half-lives beyond 24 h in 10% MS. While it is difficult to relate half-lives determined in vitro to in vivo experiments, the observed stabilization effect potentially offers a path towards novel therapeutic applications. By comparison, coating with decalysine-PEG_5kDa_ copolymers also stabilized **PB84** for more than 24 h under the same conditions **(Figure S5)**. Yet, these cationic copolymers bear the risk of collapsing two-helix wireframe architectures, as well as reducing antigenic activity due to steric exclusion. We further note that crosslinking with glutaraldehyde resulted in substantial band shifts upon incubation in 10% MS in our hands, likely due to residual reactivity and conjugation to serum proteins **(Figure S5)**.

We expect that MGB-based protection can further be improved by exploring additional derivatives of **5**. The level of protection conveyed by MGBs **1** to **5** correlates with their affinity for the minor groove of dsDNA and their specificity profiles. PCs **1** and **2** have the lowest affinities (apparent K_D_ = 10-100 nM)^50^ and additionally display secondary binding modes along the phosphate backbone that likely don’t compete with DNAse I binding^44-46^. While the affinities of benzamidine **3**^51^ and diamidine **4**^52^ for AT tracts is comparable to that of **5** (apparent K_D_ = 1-10 nM)^37, 50^, both MGBs have more restrictive specificity profiles and might therefore leave parts of **PB84** unprotected. The fact that stabilization against 10% MS by **5** was titratable further supports an essential role of sequence-specific affinities where low affinity sequence motifs are only saturated at higher concentrations. Extensive medicinal chemistry efforts over the past decades have produced a plethora of high-affinity MGBs with varying specificity profiles, and will be explored in future studies^39^. Here, the use of combinations of MGBs matching the sequence space of a given DNA nanoparticle seems promising. Our initial investigations combining **5** with MGBs **1, 3** or **4** did, however, not result in improved half-lives in 10% MS **(Figure S5)**.

Next, we explored whether diamidine-based stabilization was broadly applicable to other geometries. Coating of a two-helix icosahedron (**I52**) with **5** also resulted in substantial protection against 10% MS and half-lives beyond 12 h **(Figure 4a)**. Interestingly, the overall stability of this geometry was lower than that of **PB84**, despite denser nanostructuring and higher rigidity due to a higher vertex- and edge-to-base pair ratio. Notably, concentrations were normalized per base pair and the DNA nanoparticle concentration of the larger **I52** were thus lower. This has been previously described to substantially affect the results of degradation assays and might influence our observations^53^. Finally, to demonstrate the utility of our stabilization strategy for translational applications, we investigated its compatibility with DNA origami functionalization. Toward this end, we synthesized **PB84** conjugated to 10 copies of the engineered HIV antigen eOD-GT8, using PNA:DNA hybridization, as previously described^14^. The functionalized two-helix wireframe assembly was subsequently coated with diamidine **5** and incubated with Ramos B cells recombinantly expressing an IgM-BCR specific for eOD-GT8^54-55^. B-cell activation measured by Ca^2+^ flux was comparable for non-protected and protected DNA nanoparticles **(Figure 4b)**. These findings highlight that diamidine-based stabilization can be implemented post-functionalization without disruption of structural integrity, and may therefore be readily compatible with most therapeutically relevant bioactive molecules. And because MGBs do not rely on steric exclusion, but rather on competitive inhibition, with their low molecular weight ensuring accessibility of bioactive molecules presented by DNA origami, they offer a valuable approach to stabilizing DNA origami both in vitro and in vivo.

**Figure 4.**
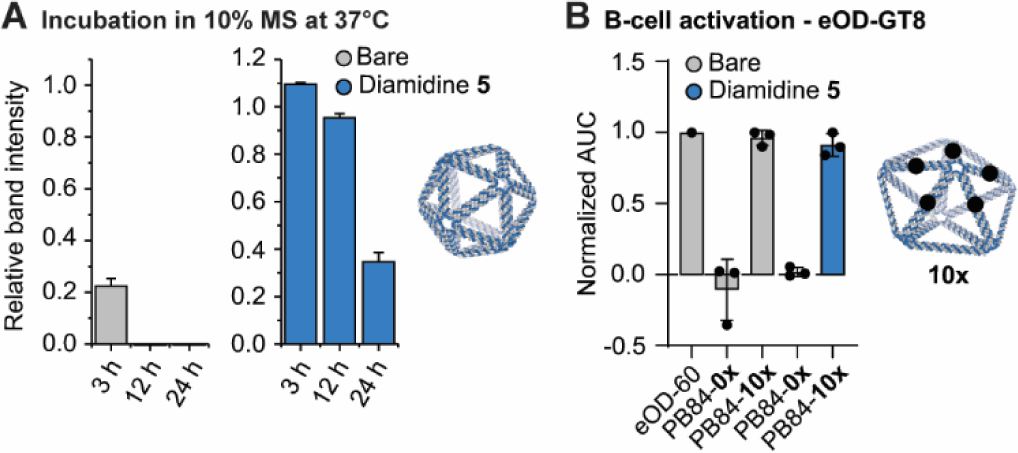
Applicability of diamidine-based stabilization to different wireframe architectures and functionalized DNA origami. **(A)** Coating with diamidine **5** also conveyed protection against 10%MS to **I52**. Relative band intensities compared to the 0 h data point for bare and coated DNA nanoparticles are shown. **(B)** Diamidine **5**-coating of **PB84** functionalized with 10 copies of the engineered HIV antigen eOD-GT8 did not affect Ramos B-cell activation *in vitro*. A previously published protein nanoparticle presenting 60 copies of eOD-GT8 (eOD-60) served as the positive control for the Ca^2+^ flux assay. The calcium flux assay was conducted at total eOD-GT8 concentrations of 5 nM. All experiments were performed in triplicate. Representative gel images are shown.

## Conclusions

We introduce a facile, scalable and low-cost stabilization strategy for wireframe DNA origami using MGBs. While unprotected two-helix wireframe architectures were found to be stable for at least 24 h under cell culture conditions, they degraded rapidly at physiological DNAse I concentrations and in MS. Coating of **PB84** with different MGBs conveyed substantial protection under both conditions via competitive inhibition of endonuclease activity. Here, diamidine **5**, also known as DAPI, was identified as a particularly potent stabilizer extending half-lives beyond 24 h, likely due to its high affinity and broad sequence specificity. We further demonstrate that our strategy is broadly applicable to varying geometries and compatible with post-folding DNA origami functionalization. Due to their binding mode, diamidines convey lower positive charge densities to DNA origami compared with established cationic polymer-based methods. Therefore, MGB coating represents a viable, complementary stabilization approach for preclinical studies and therapeutic indications.

While our findings represent a proof-of-principle for MGB-based protection of virus-like DNA origami, several technical challenges remain and need to be addressed in future studies. On the one hand, our findings indicate that the extent of DNA origami protection against nucleases is limited by the coverage of diverse sequence motifs. It will therefore be important to identify MGBs or combinations of MGBs that display both low dissociation rates and broad sequence specificity. Particularly in vivo, where DNA nanoparticles and dissociating MGBs are highly diluted upon injection, stabilization efficiency potentially decreases over time. The use of MGB-alkylating agents conjugates represents an attractive strategy to address this challenge via the formation of covalent bonds. Moreover, we have not yet explored the third canonical class of MGBs, pyrrole-imidazole polyamides. For this class of compounds, recent advances in synthetic methodology and the development of hairpin structures facilitate the tuning of sequence specificity and leverage avidity effects^42-43^. This might enable the rational design of MGBs tailored to the sequence space of a given DNA nanoparticle.

On the other hand, MGBs are chemotherapeutics with clinical relevance for the treatment of infectious diseases and cancer. While this also offers opportunities to improve the challenging delivery of MGBs via DNA origami in vivo, the compounds might also be inherently toxic. Here, dosing studies in mice and the identification of potent stabilizers with favorable toxicity profiles will be essential. Importantly, the FDA approval of furamidine as an experimental drug and the fact that several other MGBs have been under clinical investigation, indicate that compounds with these properties can, in principle, be designed for a given therapeutic indication (NCT 03824795 is an ongoing phase II clinical trials)^40-41^. We thus envision the parallel exploration of this dual utility of MGBs in the context of DNA origami, either as biocompatible stabilizers or as therapeutic cargo.

## Methods

Methods are described in the **Supporting Information**.

## Acknowledgments

We thank Xiao Wang for synthesizing the HEG-modified oligonucleotide staples.

This work was supported by the Human Frontier Science Program (RGP0029/2014), the Office of Naval Research (N00014-16-1-2953), the U. S. Army Research Office through the Institute for Soldier Nanotechnologies at MIT (Cooperative Agreement Number W911NF-18-2-0048), the Ragon Institute of MGH, MIT, and Harvard, the Marble Center for Nanomedicine, and the NIH (R21-EB026008, R01-MH112694, AI048240, UM1AI144462, and UM1AI100663). E.-C. W. is supported by the Feodor Lynen Fellowship of the Alexander von Humboldt Foundation. DJI is an investigator of the Howard Hughes Medical Institute.

