## Supporting Information for "Controlling wireframe DNA origami nuclease degradation with minor groove binders"

#### Title

#### Methods

##### Materials

SS320 E. coli cells were purchased from Lucigen. EndoFree GigaPrep Kits were purchased from Qiagen. Tris acetate-EDTA (TAE), Tris borate-EDTA (TBE) and PBS buffer were purchased from Corning. The phosphate clamps **1** and **2** were prepared as previously described[1]. The minor groove binders (MGBs) **3** (pibenzimol), **4** (furamidine) and **5** (DAPI, 0.1% (w/v) in $H_2O$) were purchased from Sigma Aldrich. PEGylated oligolysine ($K_{10}$-$PEG_{5kDa}$) was purchased from Alamada Polymers. Oligonucleotide staples (purified via desalting columns) were ordered from IDT. The standard base and hexaethylene glycol (HEG) phosphoramidites were purchased from Glen Research. Agarose was purchased from IBI Scientific. $MgCl_2$, NaCl, $PEG_{8kDa}$, antibiotics, 2x YT medium, TE buffer, ethidium bromide (EtBr), glutaraldehyde (50% (v/v) in $H_2O$), TritonX-144 and Pluronic F-127 (10% (v/v) in $H_2O$) were purchased from Sigma-Aldrich. DMEM, FBS and MS were purchased from Gibco. DNase I and Exo I were purchased from New England Biolabs. Amicon Ultra centrifugal filters (100 kDa) were purchased from Sigma Aldrich. Zeba spin columns (7 kDa) and Proteinase K were purchased from ThermoFisher.

**Scaffold synthesis**

The custom-length DNA scaffold (phPB84, 2520 nt) for **PB84** was prepared as previously described[2]. The custom-length DNA scaffold (phI52, 3210 nt) for **I52** was prepared following an adapted protocol based on this general approach. Briefly, the required plasmid was generated by extending the synthetic sequence of the phPB84 plasmid by Gibson assembly, with the additional synthetic sequence amplified from pUC19 vector. Next, SS320 E coli cells were transformed with the phI52 plasmid and the M13cp helper plasmid (provided by Dr. Andrew Bradbury, Los Alamos National Laboratories). After clone selection, sequence identity validation and time course optimization for the selected clone, with all steps using 2x YT medium containing 100 µg/mL ampicillin, 15µg/mL chloramphenicol and 5µg/mL of tetracycline, transformed cells were used for phI52 production for DNA origami folding. Pre-cultures were grown overnight at 37°C, diluted 100-fold and incubated for another 8 h, with all steps using 2x YT medium containing 100 µg/mL ampicillin, 15µg/mL chloramphenicol and 5µg/mL of tetracycline. Cells were sedimented by centrifuging three times at 4000 g for 3 min and subsequently discarded. Phage was precipitated from the supernatant in presence of 6% (w/v) and 3% (w/v) of NaCl by stirring at 4°C for 1 hour and harvested by centrifugation at 20,000 g at 4°C for 1 h. After resuspension if TE buffer, ssDNA was extracted via the EndoFree GigaPrep purification protocol with the following modifications: Proteinase K was added to buffer P1 followed by incubation at 37°C for 1 h, addition of buffer P2 and incubation at 70°C for 10 minutes. After ssDNA purification, Triton X-144 was used to improve endotoxin removal[3]. Purity of the scaffold was analyzed by agarose gel electrophoresis (AGE) (1.6% agarose, TAE buffer with 12 mM $MgCl_2$, EtBr, 65V for 150 in at 4°C) and sequence identity was validated by primer walking with Sanger sequencing **(Figure S1, Tables S1 and S4)**.

**Oligonucleotide staple synthesis**

Solid-phase DNA synthesis was performed on a Dr. Oligo synthesizer purchased from Biolytic. DNA synthesis was performed on a 200 nmol scale, starting from universal 1000 Å CPG solid-supports and following the standard protocol[4]. Standard base and HEG phosphoramidites were dissolved in anhydrous dichloromethane to afford 0.1 M solutions and were used in 10-fold excess at a coupling time of 1 min. Coupling efficiency was monitored after removal of the dimethoxy trityl (DMT) 5″-OH protecting groups. After solid-phase synthesis, oligonucleotides were fully deprotected in concentrated ammonium hydroxide at 60°C for 3 h and purified over desalting columns. Installation of the HEG modification was validated by denaturing polyacrylamide gel electrophoresis (PAGE) (8M urea and 20% PA, TBE buffer, SybrGold, 120V for 180 min at 25°C) of **PB84** oligonucleotide staple mixtures **(Figure S2)**.

**DNA nanoparticle design and assembly**

**PB84** and **I52** were designed using DAEDALUS and nick position of edge staples were adjusted manually for outward orientation[5] **(Table S2 and S3)**. DNA nanoparticles were assembled as previously described[5]. Briefly, Briefly, 30 nM of scaffold and 300 nM of each oligonucleotide staple were dissolved in TAE buffer with 12 mM $MgCl_2$ and thermally annealed as follows: 95°C for 5 min, 80–75°C at 1°C per 5 min, 75–30°C at 1°C per 15 min, and 30–25°C at 1°C per 10 min. HEGylated **PB84** was assembled exclusively from staples that were 3' and 5' modified. In case of **PB84-10x**, bearing 10 overhang sequences (TTACTGGACTG), a reverse-complement to this sequence was added to the reaction mixture at 2-fold excess. **PB84** and **I52** were purified into PBS using Amicon Ultra centrifugal filters (100 kDa, 2000 g, 3x 15 min) and stored at 4°C. **PB84-10x** was purified into TAE buffer with 12 mM $MgCl_2$ using Amicon Ultra centrifugal filters (100 kDa, 2000 g, 3x 15 min) and stored at 4°C. Purity and monodispersity of the DNA nanoparticles were validated by AGE (1.6% agarose, TAE buffer with 12 mM $MgCl_2$, EtBr, 65V for 150 in at 4°C) **(Figure S1 and S2)**.

**DNA nanoparticle coating**

For all MGBs and PEGylated oligolysine ($K_{10}$-$PEG_{5kDa}$), 1 mM stocks in $H_2O$ were prepared. Prior to use degradation or B-cell activation assays, 15 nM **PB84** or 12 nM **I52** were incubated with 40 µM MGBs in DMEM for at least 30 min at room temperature, corresponding to an MGB:base pair ratio of 1. Mixtures of MGBs were prepared so that the total MGB concentration corresponded to 40 µM. Coating with PEGylated oligolysine and glutaraldehyde crosslinking was performed as previously described[6]. Briefly, 180 nM **PB84** were incubated with 90 µM PEGylated oligolysine in PBS for at least 60 min at room temperature, corresponding to an amino group:base pair ratio of 1. For glutaraldehyde crosslinking, 90 nM coated **PB84** was incubated with 2% (v/v) glutaraldehyde for 120 min at room temperature. The DNA nanoparticles were subsequently purified into PBS using Zeba spin columns (7 kDa).

**Degradation assays – serum**

15 nM **PB84** or 12 nM **I52** were incubated in freshly prepared 10% FBS or MS in DMEM at 37°C at a total volume of 30 ul per aliquot. Following incubation for a given time point, samples were cooled to 4°C and analyzed by AGE (1.6% agarose for bare and MGB-coated and 1.2% agarose for PEGylated oligolysine-coated DNA nanoparticles, TAE buffer with 12 mM $MgCl_2$, EtBr, 65V for 150 min at 4°C). Additionally, denaturing AGE (addition of 2 mM EDTA and incubation in 50% formamide at 37°C for 30 min prior to loading, 1.6% agarose, TAE buffer, EtBr, 65V for 150 min at 4°C) and denaturing PAGE (addition of 2 mM EDTA and incubation in 50% formamide at 37°C for 30 min prior to loading, 8M urea and 20% PA, TBE buffer, SybrGold, 120V for 180 min at 25°C) were performed for bare **PB84** in 10% FBS. For PEGylated oligolysine-coated DNA nanoparticle, 5-fold excess of PEGylated oligolysine was added to the sample prior to AGE to ensure migration into the gel.

**Degradation assays – nucleases**

15 nM **PB84** were incubated with 0.5 U/ml DNAse I or Exo I in the respective nuclease-specific reaction buffer at 37°C at a total volume of 30 ul per aliquot. Following incubation for a given time point, samples were cooled to 4°C and analyzed by AGE (1.6% agarose for bare and MGB-coated, TAE buffer with 12 mM $MgCl_2$, EtBr, 65V for 150 in at 4°C).

**DNA nanoparticle functionalization and B-cell activation assay**

**PB84-10x** was functionalized with PNA-modified (-GGK-cagtccagt-K) eOD-GT8 as previously described[7-8]. Briefly, 50 nM of **PB84-10x** were incubated with 2.5 µM PNA-modified eOD-GT8 in PBS and thermally annealed as follows: 37°C to 25°C at 1°C per 20 min. Functionalized **PB84-10x** was purified into PBS using Pluronic F-127-coated Amicon Ultra centrifugal filters (100 kDa, 2000 g, 3x 15 min) and stored at 4°C[9]. Purity and monodispersity of the DNA nanoparticles were validated by AGE (1.6% agarose, TAE buffer with 12 mM $MgCl_2$, EtBr, 65V for 150 in at 4°C). Quantitative coverage (>90%) with eOD-GT8 was validated via tryptophan fluorescence. The characterization of the used DNA nanoparticle sample was published previously[7]. 30 nM functionalized **PB84-10x** were coated at 80 µM of diamidine **5** in PBS. Quantitative coverage (>90%) with eOD-GT8 was validated via tryptophan fluorescence.

The B-cell activation assay was conducted as previously described[7]. Briefly, human Ramos B cells, stably expressing the VRC01 germline IgM B cell receptor[10], at a cell density of 10 million cells/ml were incubated with 10 µM Fluo-4 AM for 30 minutes at 37°C. After washing, calcium flux assays were performed on plate reader at 37°C on a 96 well microplate with 160 µL of Fluo-4 AMlabeled human Ramos B cells at 2 million cells/ml. The baseline fluorescence at 505 nm was recorded for 1 min. Next, 40 µL of nanoparticles were added to the cells to afford a final eOD-GT8 concentration of 5 nM of antigen and fluorescence measurements were continued for another 7 min. After baseline subtraction, fluorescence traces were integrated and normalized using the integrated fluorescence intensity obtain for the eOD-60mer to obtain the normalized AUC. Student's t-test (n = 3, α = 0.05) was performed to test for effects of diamidine **5** on B-cell activation.

#### Supporting Figures

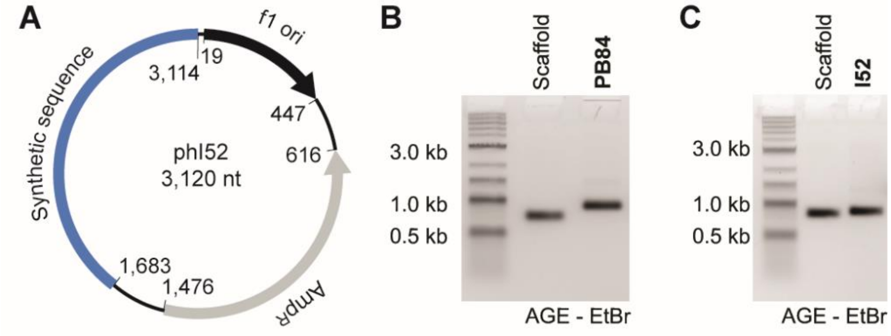


**Figure S1. Phagemid map and characterization of DNA nanoparticle synthesis.**
**(A)** The phagemid map for the custom-length, circular scaffold (phI52, 3210 nt) used to synthesize **I52** is shown with features and corresponding nucleotide positions highlighted. **(B)** The characterization of the used phPB84 scaffold and purified **PB84** by AGE is shown. **(C)** The characterization of the used phI52 scaffold and purified **I52** by AGE is shown.

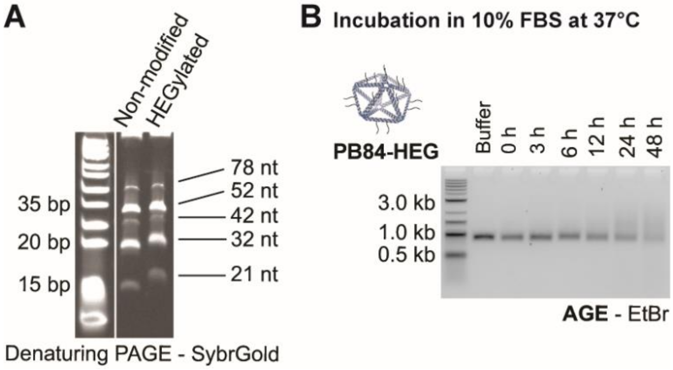


**Figure S2. Stability of HEGylated PB84 in 10% FBS.**
(**A**) Characterization of synthesized oligonucleotide staples for **PB84-HEG** by denaturing PAGE. The comparison between the non-modified and HEGylated staple mixes reveals band shifts for all oligonucleotide staples. **(B)** Incubation under typical cell culture conditions, in DMEM with 10% FBS revealed that **PB84-HEG** was not stabilized compared to non-modified **PB84.** Degradation experiments were conducted in triplicate. Representative gels images are shown.

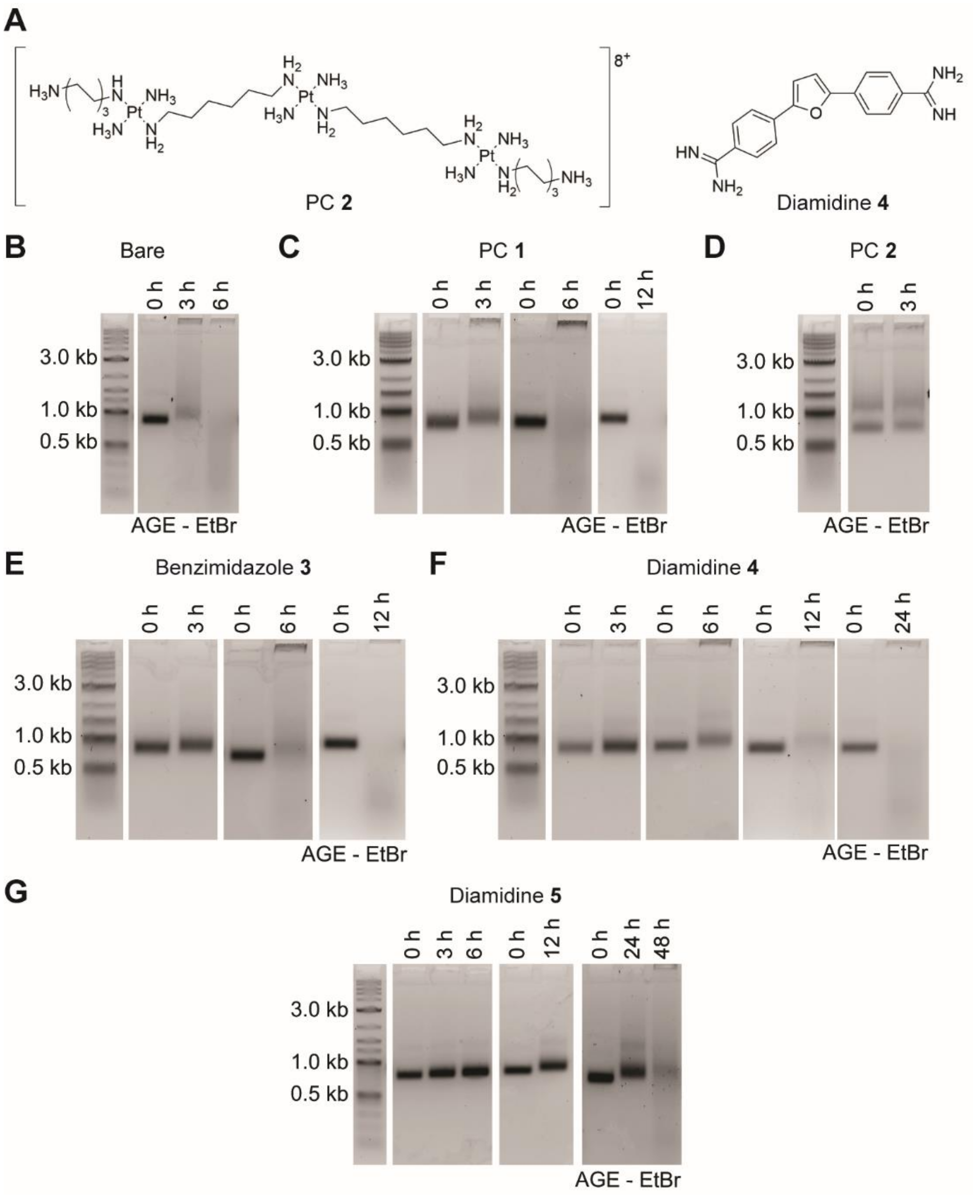


**Figure S3. Supporting information for stabilization of PB84 with MGBs.**

**(A)** The structures of PC **2** and diamidine **4** are shown. **(B to F)** Representative gel images of the screening of MGBs **1** to **5** as stabilizers of **PB84** for incubation in 10% MS at 37°C are shown. All experiments were conducted in triplicate.

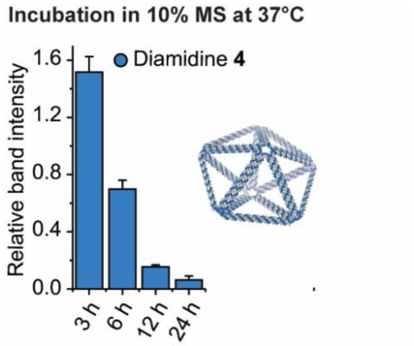


#### Figure S4. Stabilization of PB84 with diamidine 4.

The quantification of degradation of **PB84** coated with diamidine **4** in DMEM with 10% MS at 37°C revealed substantial stabilization compared to bare DNA nanoparticles. Relative band intensities compared to the 0 h data point are shown. All experiments were performed in triplicate.

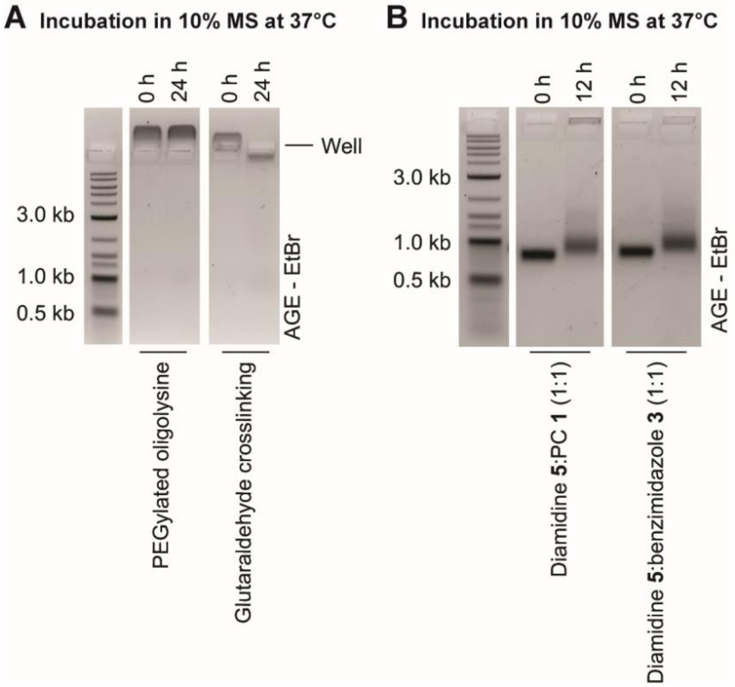


#### Figure S5. Stabilization of PB84 with PEGylated oligolysine and diamidine 5 mixtures.

**(A**) Coating of PB84 with PEGylated oligolysine as previously described[3, 6] resulted in substantial stabilization of the DNA nanoparticle for at least 24 h. While the band intensity also remained constant for coated **PB84** subsequently treated with glutaraldehyde, we observed band shifts upon incubation in 10% MS, suggesting reactivity towards serum proteins. **(B)** The coating with mixtures of diamidine **5** with PC **1** or benzimidazole **3** did not enhance stabilization compared to **5** alone (**Figure S3**). All experiments were conducted in triplicate. Representative gel images are shown.

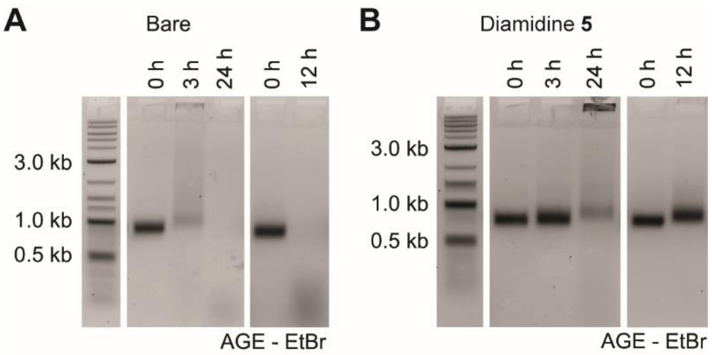


**Figure S6. Supporting information for stabilization of I52 with diamidine 5.**
(**A**) Representative gel images of the evaluation of **I52** stability in 10% MS at 37°C are shown. **(B)** Representative gel images of the evaluation diamidine **5** as a stabilizer of **I52** are shown. All experiments were conducted in triplicate.

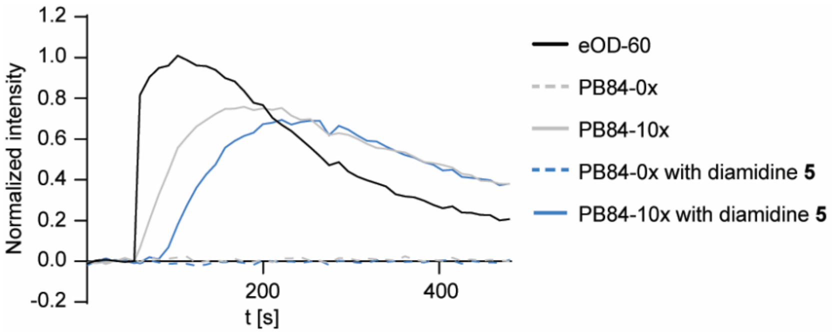


**Figure S7. Supporting information for B-cell activation assay.**
Representative fluorescence traces of human Ramos B cells loaded with Fluo-4 AM and incubated with functionalized DNA nanoparticles. (representative individual calcium trace). All experiments were conducted in triplicate (as biological replicates).

### Supporting Tables

**Table S1 – Scaffold Sequences**

**Table S2 – Staple Sequences – PB84 and PB84-10x**

**Table S3 – Staple Sequences – I52**

**Table S4 – Primer Sequences**
